# A systematic review of European farmer and non-farmer attitudes towards landscapes, ecosystem services, and agricultural management practices: Implications for permanent grassland management

**DOI:** 10.1101/2020.06.12.148585

**Authors:** Sophie J. Tindale, John Elliott, Marjolein Elings, Rosa Gallardo-Cobos, Erik Hunter, Eva Lieberherr, Simona Miškolci, Paul Newell Price, Simone Quatrini, Pedro Sánchez-Zamora, Hannah Schlueter, Lynn J. Frewer

## Abstract

Permanent grassland (PG) is an important agricultural land use for the delivery of multiple ecosystem services (ES), including carbon sequestration, water quality protection, food production, habitat provision, and cultural activities. However, PG environments are threatened by sub-optimal management, cultivation, and abandonment that are influenced by context, land manager’s attitudes and societal demand for ES. Therefore, the perceptions and attitudes of key decision-makers (farmers) and other stakeholders (non-farmers, including citizens and consumers of the products of permanent grasslands, and ES) need to be understood to ensure the sustainability of PGs and the ES they provide. A systematic review of the literature identified 135 scholarly articles. Application of thematic analysis, allowed the organization, and synthesis of current research related to (different) stakeholder attitudes, and how these influence PG management and the delivery of ES. The results suggest that different stakeholders hold different views towards permanent grassland, with farmers in particular having to balance economic with other (potentially conflicting) drivers. The types of knowledge held by different groups of stakeholders, access to sources of information, as well as the influence of knowledge on behaviour; and environmental values (for example in relation to aesthetics or conservation of biodiversity) explained why certain motivations for attitudes and behaviours are held. A major gap, however, was identified in relation to PG as opposed to other types of landscape.

## 1. Introduction

The existence and management of permanent grasslands (PG) is key to the delivery of multiple ecosystem services (ES) in Europe and elsewhere, including carbon sequestration, water protection, food production and cultural activities [1-4]. Consequently, the preservation and sustainable management of grassland is important for delivery of ES and for shaping landscapes [5]. However, PG maintenance and functions are under threat from sub-optimal management of inputs, cultivation in higher output farming systems and abandonment in remote and marginal areas [6]. The decisions made by land managers and farmers in relation to changing management options, or choices to convert or abandon PG will influence the balance and extent of ES are delivered by PG [7]. Multiple incentives and barriers linked to the wider economic system, political influences, social norms, biophysical and climatic contexts, and individual and collective behaviours underpin these decisions. The attitudes and perceptions of farmers and land managers may affect their priorities and goals of action, inaction and resultant landscape change [8-11]. Consumers and citizens, mostly non-farmers, can influence change through their willingness to use the landscape for recreation, influencing policy makers, facilitating biodiversity protection, or engaging with local businesses, or changing demands for products from grassland landscapes and livestock raised on these [12-14].

An increasing number of assessments of land management decision-making and priority setting focus on stakeholder perceptions and attitudes, either in order to understand the ES that are most relevant, or the evaluation of suitable management options [15]. Receptiveness to particular contexts, inputs, knowledge and stimuli are often moderated by existing beliefs, attitudes, motivations, and personality [16, 17]. ES have different meanings for different stakeholder groups [18] depending on their knowledge, access to networks, professional experience, cultural context and socio-economic situations [19-21]. For example, conservationists are likely to possess knowledge of the value of habitats for species protection [22-24] whereas farmers understand how to influence agricultural output, and the value of production[18]. Equally, stakeholders may ascribe different values to ES at different local, regional or institutional levels [25] based on their cultural background as well as the impact that any particular ES might have on their health and well-being [26]. Individuals can also represent different stakeholder groups at different times and in different contexts. Differences in perceptions offer insights on decisions that shape spatio-temporal trade-offs between ES [27]. Trade-offs between agricultural productivity and other ES delivery are important in relation to sustainability issues [11, 28].

Studying attitudes towards the ES that emerge from PG landscapes in different socio-cultural contexts facilitates understanding of individual preferences [29, 30] that underpin conflicts between stakeholders with diverse values, interests, experiences and knowledge [29]. To understand conflicts requires insights from the perspective of land managers (e.g. farmers) who make decisions that influence ES provision, and from those who benefit or are deprived in some way by their reduction, removal, or replacement [31]. This has consequences for the use of different stakeholders’ knowledge and preferences in the assessment, and prioritisation, of ES in grassland environments [18, 32, 33]. Failure to consider attitudes about ES, landscape preference and management practices may lead to sub-optimal management practices (Hein *et al*., 2006), which do not align with societal priorities. It is therefore important to understand the range of views and how these are traded off in relation to ES in order to inform policies and practices that lead to more sustainable land management.

There are only a small number of empirically based studies investigating farmer attitudes, behaviour and decision making around land management and ES [18] particularly in relation to grassland landscapes. Notable exceptions include [19] who investigated farmers’ perceptions and values related to ES in the Central French Alps with the focus on mountain grasslands management and Lewan and Söderqvist’s [34] analysis of how farmers in a Swedish river valley identify and value different ES. Other studies such as those by Switek and Sawinska [35], Schulz *et al*. [36] and van Herzele *et al*. [37] conducted surveys looking at the importance of greening activities and how perceptions around the related ES might affect agricultural management practices. Similarly, Sattler and Nagel [38] investigated the level at which farmers will accept and adopt conservation measures.

Societal attitudes towards land management have tended to focus on specific issues such as conservation, aesthetics, stakeholder well-being, climate change and recreation, particularly within the theme of ES [39-41]. Other studies have contrasted opinions of farmers with interested non-farming populations [42, 43].

Although research on these topics is not new, the scope is broad. To the best of our knowledge no systematic review has integrated knowledge on the perceptions and attitudes of stakeholders to landscapes, ES and management practices in the context of PG in Europe. Understanding how different societal actors, including farming and non-farming stakeholders perceive landscapes and the associated ES, is a significant challenge for better understanding land management decisions and societal impacts, and their changes over time [44]. This is particularly important in relation to the link between perceptions and attitudes, individual intentions and behaviour, and collective social, economic and political decision-making [45].

To address these gaps, a systematic literature review was conducted to organize, synthesize and discuss the current knowledge base related to stakeholder attitudes and how they influence PG management and the delivery of ES across Europe. Specifically, two key questions relating to stakeholder attitudes were identified: i) What are the attitudes of different stakeholder groups (including farmers and non-farmers) to grasslands, grassland landscapes including PG, land management in PG environments and ES delivery? ii) What factors influence the perceptions and attitudes of different stakeholders towards PG and ES? The aim was to show how (differences in) attitudes can subsequently be considered in the development process of alternative management scenarios [46] and in the assessment of ecosystem resilience and socio-economic implications of such land-use change decisions; and to inform policy-making processes, e.g. for the establishment of incentive, payments or compensation mechanisms [47-50]. Gaps in knowledge were identified, to facilitate the design of future research into perceptions of, and attitudes towards, PG and ES.

The studies included encompass those that are grassland specific as well as those that are more general, such as agro-ecological land use and management, where grassland is a part of the system. For this reason, the review incorporates grassland and other wider landscape systems that include PG, as PG is not always referred to as such in the published literature.

## 2. Materials and Methods

### 2.1. Literature search

A systematic review [51, 52] protocol was developed (see S3 Protocol) to meet the study objectives and to address the groups of interest, specific conditions, comparisons, outcomes and the context of the study. Search terms were derived from the initial research questions using appropriate key words. Search strings were trialled and refined in a multistep process, with the face validity of each search addressed by assessing the appropriateness of the papers returned, and by checking some search results for key authors identified through an initial search. A wide variety of search terms were used in order to cover the multiple contexts in which PG could be perceived and in which decisions could be made. All strings included phrases aimed to find perceptions, attitudes, opinions, values, preferences and choices. Additional specific topic words or phrases were added to each search. Two examples of search strings are shown below:

> “permanent grassland” AND (attitude* OR opinio* OR willingness OR accept* OR prefer* OR percept* OR belief OR trust OR valu*)
>
> “rural landscape” AND “grassland” AND “citizen” AND (attitude* OR opinio* OR willingness OR accept* OR prefer* OR percept* OR belief OR trust OR valu*)

Overall, 11 search strings were used (see Appendix 1). The search was conducted on the 5th and 6th of February 2019, with some additional searches added on 11th and 14th April 2019. To increase the likelihood of discovery and reduce bias in the algorithms used in different search engines, all searches were repeated in Scopus and Google Scholar [53].

Reporting followed the Preferred Reporting Items for Systematic Reviews (PRISMA) [54] (Figure 1). Initially the search returned 96,877 records. Where there were more than 100 results in both Google Scholar or Scopus searches, only the first ∼100 records were downloaded and included in the review (equivalent to 10 pages of Google Scholar results), thus capturing the most relevant hits while still being a feasible amount to screen [55]. The searches resulted in 2392 documents included for screening after the removal of duplicates.

**Fig1.**
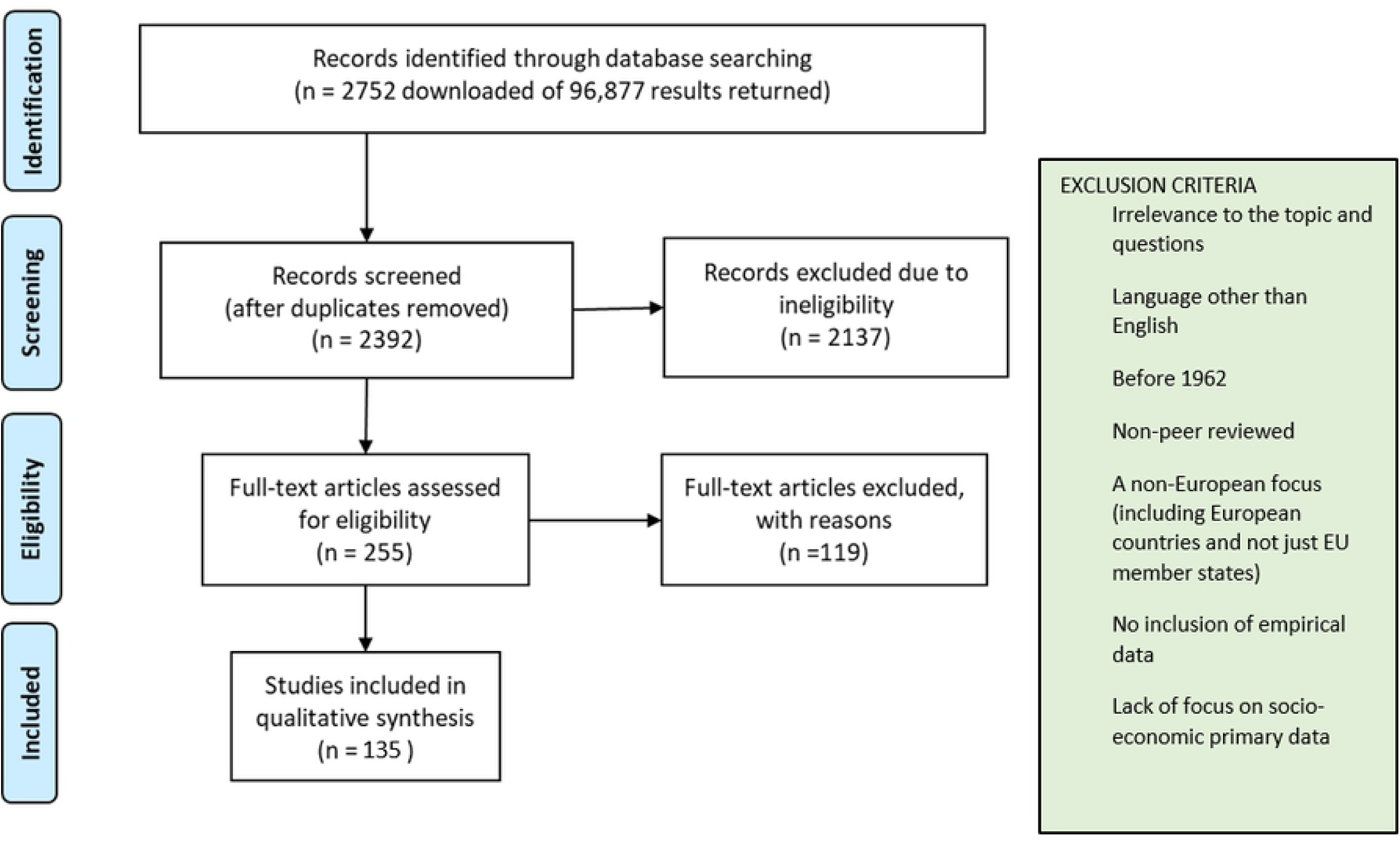
PRISMA flow diagram of systematic review process and exclusion criteria used.

### 2.2. Selection and screening of relevant studies

Peer reviewed journal articles were screened using a pre-defined set of exclusion criteria (see Figure 1). Papers before 1962 were excluded as this was before the advent of the EU Common Agricultural Policy (CAP), which potentially has had very significant influence on land management and the economic and social context of farming, and therefore on perceptions and decision-making in relation to farming practice. Papers included were those that focused on empirical data collection (therefore excluding literature reviews and theoretical pieces). In addition, the empirical data were required to be socio-economic primary data, although hybrid studies of physical science data (e.g. climate data) and social data were included.

The results of the literature search were exported to a Mendeley library, and duplicates were removed. Studies were then screened, in a two-stage process using a traffic light system derived and verified amongst the research team (green = included; amber = uncertain, another team member to review; and red = excluded). Ten % of the studies were cross checked for agreement in coding. First, the titles and abstracts were screened for relevance. Then the resultant (255) articles were read in full. This process allowed the relevant articles to be analysed for themes and information, and to be assessed against critical appraisal criteria (e.g. [56]), and to allow further screen for irrelevant studies. One hundred and thirty-five articles were eventually included in the thematic analysis.

### 2.3. Thematic Analysis

Full text articles were categorised and analysed using QSR Nvivo 12. For each article, basic information was recorded including: location of the study (country), research problem or premise, target group (e.g. livestock farmers), aims and objectives (the core purpose of the study), date or timespan of the study (date when the data were collected), methods used (e.g. survey), and any critical appraisal issues.

Three broad themes were used to categorise data relating to the aims of the systematic review: ‘Attitudes and perceptions’, ‘factors affecting attitudes’, and ‘factors affecting decision-making’. Open coding was used to derive sub-themes [57, 58] and focused coding was conducted to break down themes further into sub-categories to aid explanation and exemplification of the intricacies of the original codes [59].

In relation to ‘attitudes and perceptions’, eight sub-themes were identified, each of which contained up to 19 sub-categories, and up to 15 sub-sub-categories (Table 1). A similar pattern emerged for ‘factors affecting attitudes’, where 47 sub-categories were identified (again, some with sub-nodes themselves). These codes were the initial, in-depth, coding, which aimed to capture the depth and breadth of information within the studies. Codes were gradually assimilated into more coherent themes through a process of analysis.

**Table 1.** Themes and examples of sub-themes coded in Nvivo 12 from 135 full text articles for the categories of ‘attitudes and perceptions’ and ‘factors affecting attitudes’.

| Theme | Examples of sub-themes |
| --- | --- |
| Attitudes to ecosystem services (ES) | *Cultural ES; decision-makers perceptions; farmers perceptions; gaps, conflicts and trade-offs; multiple ES; citizens or residents perceptions of ES; scientists perceptions.* |
| Attitudes to grassland | *Discourses; ecosystem services; grassland abandonment; grazing and grazing systems; soil fertility; species richness.* |
| Attitudes to the environment and farmland | *Abandonment; biodiversity, care for the land; conflict or variability of attitudes; food production; multi-functionality; rewilding and wilderness; towards farmland and landscape; towards conservation; towards national parks; towards nature; towards traditional landscape.* |
| Attitudes to management practices | *Agroforestry; impact of management practices; agri-environment schemes; innovation and technology; organic; policy; risk; variability of attitudes.* |
| Attitudes of consumers and the public | *Agriculture; grazing; landscape; local community; transhumance.* |
| Perceptions of being a farmer | *Conservationist vs productivist; economic attitude; stewards of nature.* |
| Other themes | *Attitudes to climate change, emissions and greenhouse gases; Willingness to pay* |
| Factors affecting attitude | *Age; access to landscape; biophysical factors; complexity; education; emotions; employment status; extreme events; farm structure and system; farmer identity – farmer type or style; fulfilment; gender; geographical location and climate; historical context; income and economics; knowledge; land ownership;* |
|  | *landscape type; management; naturalness and environment; perceptions and values; political agenda; temporal and spatial scale; socio-cultural background; urban vs rural residence.* |

## 3. Results

### 3.1. Overview

In the relevant papers for review 24 European countries were represented. The UK was the most commonly studied country, with 26 papers. Studies originating in Germany (20) and Italy (20) were also numerous, as were studies focusing on multiple countries (20). Other countries covered by the papers included France (19), Austria (12), Spain (11), Denmark (9), Ireland (9), Switzerland (8), Slovenia (5), Sweden (5), Netherlands (5), Belgium (4), Slovakia, Norway, Hungary, Bulgaria, Greece (all 2), and Lithuania, Estonia, Finland, Poland and Croatia (all 1). Most studies are focused on central and northern Europe with fewer studies in Eastern Europe.

There was a wide variety of topics covered by the papers, with the most numerous focusing on adoption of agro-environment management practices (26), followed by papers focusing on different perceptions of ES (21) and those focusing on landscape preferences (13). Other common themes included attitudes to conservation of landscapes (8), farmers’ landscape decision-making (8), engagement in climate change related issues or mitigation (6), and public opinion of (future and past) rural landscape (change) (5).

Studies were published between 1990 and 2019 (Figure 2), with the most studies published in 2016 and 2018. The papers targeted both farming and non-farming stakeholders in their research and data collection (Table 2); 23% targeted both types of stakeholder, 32% targeted only non-farming stakeholders and 45% targeted only farmers (Figure 2).

**Table 2.** Types of stakeholders targeted for research and data collection including farming stakeholders and non-farming stakeholders.

| **Farming stakeholders** | **No. of papers** | **Non-Farming stakeholders** | **No. of papers** |
| --- | --- | --- | --- |
| 'Farmers' (unspecified types), | 46 | Locals/ residents, citizens, the public and non-experts | 36 |
| Livestock farmers | 10 | Other relevant stakeholders without specifying the exact mix of stakeholders | 18 |
| Dairy farmers | 8 | Government (agencies) or public officials (local and national) | 15 |
| Arable farmers | 5 | NGOs | 11 |
| Organic farmers | 4 | Agricultural extension or farm advisors | 6 |
| Conventional farmers | 3 | Distinct local populations and general or regional populations | 5 |
| Farmers implementing specific agri-environment schemes | 3 | Conservationists | 3 |
| Family farmers | 3 | Land managers | 3 |
| Upland farmers | 2 | Decision-makers | 3 |
| Farmers with more than 2ha of land | 2 | Farming organisations | 2 |
| Retired farmers | 2 | Agricultural companies | 1 |
| Peri-urban farmers | 1 | Landscape professionals | 1 |
| Farmers undertaking landscape changes | 1 | Hunters and hunting associations | 1 |
| Hobby farmers | 1 | Engineers | 1 |
| Young farmers | 1 | Professionals | 1 |

**Fig2.**
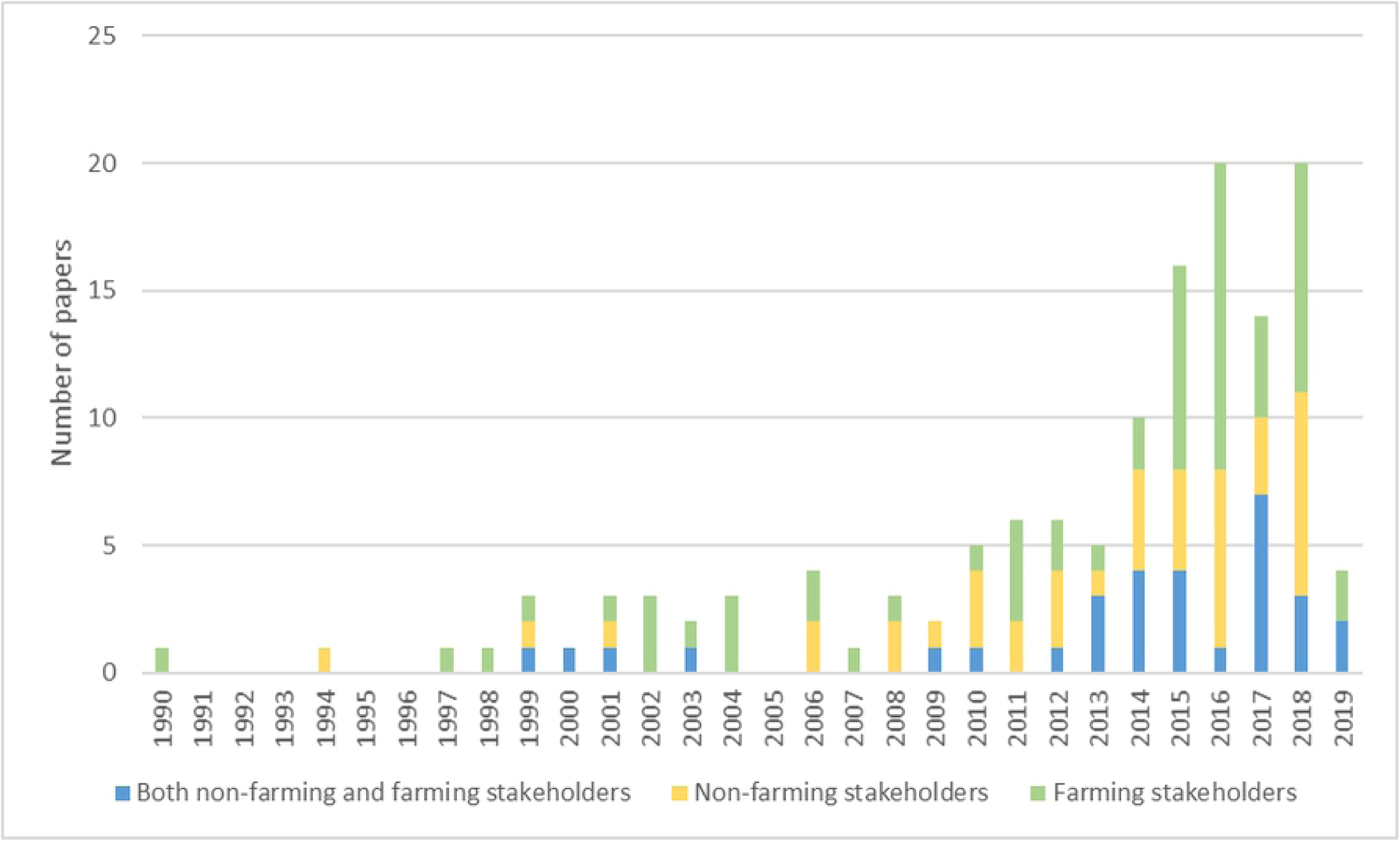
Date of publication of papers in the review including types of stakeholders targeted in research and data collection.

Data reported in studies was collected between 1988 and 2017, with many studies spanning one or two of these years. Many studies (32) do not name the date of data collection. The most common method was the survey (or questionnaire) (73); used to collect information on perceptions, behaviours and attitudes. The second most popular method was interview (47). Many studies used a combination of methods, applied in multiple stages, with interviews and surveys often being combined. Modelling was also a method applied to studies (13) in a number of ways. Participatory workshops were also used (13). Choice experiments were popular (8), particularly when trying to better understand the perceptions of participants in relation to visual landscape features (e.g. [60] – Germany), as well as payment for ecosystem services and willingness to pay (e.g. [29] – Spain).

### 3.2. Attitudes to grassland, grassland management and ES

Seventeen studies addressed attitudes to grassland and grassland management, through the study of livestock systems and case studies on specialist grassland systems such as meadows, species-rich environments, pastures and alpine grassland areas. However, it was more likely that mention of grassland was incorporated into studies of broadly defined landscapes and farming systems.

Quetier *et al*. ([41] – France) reported that the notions used by non-farming stakeholders to describe local grasslands demonstrated knowledge of, and interest in, various benefits provided by these grasslands. Cultural ES (e.g. aesthetics and testimonies to past use of the land) dominated the description of grasslands. Such recognition of benefits from grassland appeared to be more prevalent in European countries than non-European countries ([61] – multiple countries).

Multiple stakeholders saw species-rich grasslands to be an important focus of conservation efforts. This appeared to be justified for aesthetic reasons ([62] – Germany), where the public’s aesthetic appreciation was seen to increase with grassland species richness, modified by the presence of particular species. Both managed and unmanaged grassland (in the form of meadows) were appreciated by visitors and tourists for their beauty and biodiversity, and valued for their restorative health benefits ([63] – Austria). However, unmanaged meadows were seen as more interesting and offered people more to explore and discover, despite perceptions that abandoned meadows were more chaotic and confusing than the managed meadows, with a higher level of distraction. Such studies demonstrate that social benefits of particular types of grassland are not clear cut, making it complex for farmers to make decisions-about land management, particularly if demand for certain benefits is unknown. Moreover, farmers are incentivised to deliver biodiversity and landscape outcomes through voluntary engagement in agri-environment contracts, but this does not take account of the aesthetic priorities of users of the landscape, such as local residents or visitors.

Aesthetic appreciation by visitors was also found to be associated with presence or lack of grazing livestock on grasslands, depending on social factors. In a study in Germany [60], it was found that there were preferences for low numbers of grazing livestock in the landscape, in contrast to the findings in other contexts. This was attributed to the familiarity of people with the local landscape where livestock are increasingly kept indoors and are therefore less evident than in the past. The way that people perceive grassland can therefore be linked to their experience.

Other benefits provided by managed grassland included the prevention of wild forest fires. In a study in Spain [64] the prevention of forest fires was discussed by farming stakeholders in connection with five different practices related to forest and shrub clearing, and to appropriate grazing management (winter use of pastures to force animals to graze marginal areas, and use of fencing). This was an important aspect of discussion for beef farmers. They also identified that there were trade-offs associated with grazing, e.g. the optimal grazing pressure also affected other regulating services, such as “soil fertility” (natural fertilization with urine and dung), “waste management” (prevention of water pollution), and “erosion prevention” (avoidance of excessive trampling and maintenance of slopes). [64] showed that the farmers also associate the intensity of the system of grazing with the delivery of ES.

One study demonstrated that attitudes to grassland can differ between stakeholder groups. Cebrián-Piqueras *et al*. ([18] – Germany) reported a difference in preference between farmers and conservationists: conservationists preferred wet, extensively used grasslands over intensively used grasslands, and appreciated the conservation value, carbon sequestration and soil fertility of the land over forage production. Conversely, farmers preferred intensively used grasslands; soil fertility and conservation value over carbon sequestration. Another study into attitudes to grassland landscape found that locals and visitors to an area around Villar d’Arène in the central French Alps had different ways of describing grassland [41]. They found that a richer vernacular representation, focusing on the grassland resource (and associated with a long-term interaction with the local ecosystems), was understood by locals and shared little with an aesthetic appreciation of the landscape mostly found among short-term visitors.

In relation to change in grassland areas, a study in the French Alps showed that institutional constraints meant that farmers perceived an increase in grassland to be an administrative burden, and where grassland abandonment is happening there are concerns about the impact on wild flora and fauna ([65] – France). This demonstrates that negative attitudes to grassland can occur when grassland is both increasing and decreasing, depending on the circumstances of the land use change decisions.

### 3.3. Attitudes to ecosystem services

In studies referring to landscapes including grassland the concept of ES was not always understood by non-expert populations (see also [66]). Plieninger *et al*. ([67] – Germany) found that only when stressing “landscapes” rather than “ecosystems,” and “values” rather than “services” was the ecosystem-services approach useful as a unifying framework for stakeholder interaction. Often ES were inherently known and applied by farmers, whether through awareness of the term through a research project, or guessing its meaning from the word ‘ecosystem’, or via an understanding of soil function ([68] – Germany). In one study, although farmers did not demonstrate spontaneous knowledge of the ES concept, they showed an understanding of multiple ES and the agricultural practices that influenced their delivery. Bernués *et al*. ([64] – Spain) state that farmers often hold rich mental concepts of ES, even if they are not familiar with the formal terminology (Fischer and Young, 2007). Farmers also cognitively established various interactions among concepts, recognizing the complexity of ecological processes in agroecosystems ([20] – Spain).

Table 3 illustrates the key comparative findings for farming and non-farming stakeholders in relation to attitudes to ES. Findings demonstrate an often contrasting perception of importance of benefits from environments and landscapes. For example, citizens often gave a higher importance to cultural and provisioning services, including aesthetic quality, well-being, recreation, traditional and local food provision, whereas farmers gave more importance to regulating services, including water and soil regulation, as well as the provision of food and raw materials. The divergences reflect the contrasting experiences and influences on opinions and values of the differing stakeholder groups. These contrasts, although often acknowledged, are not resolved in many current approaches to land management.

**Table 3.**
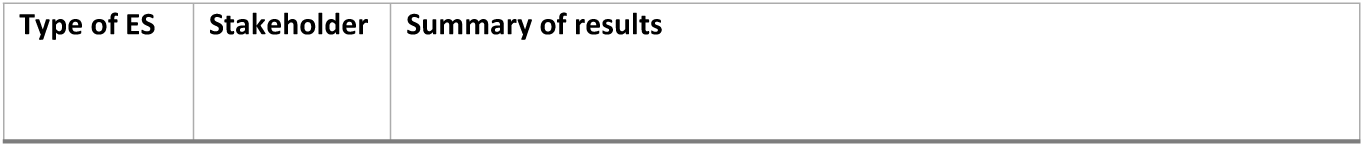
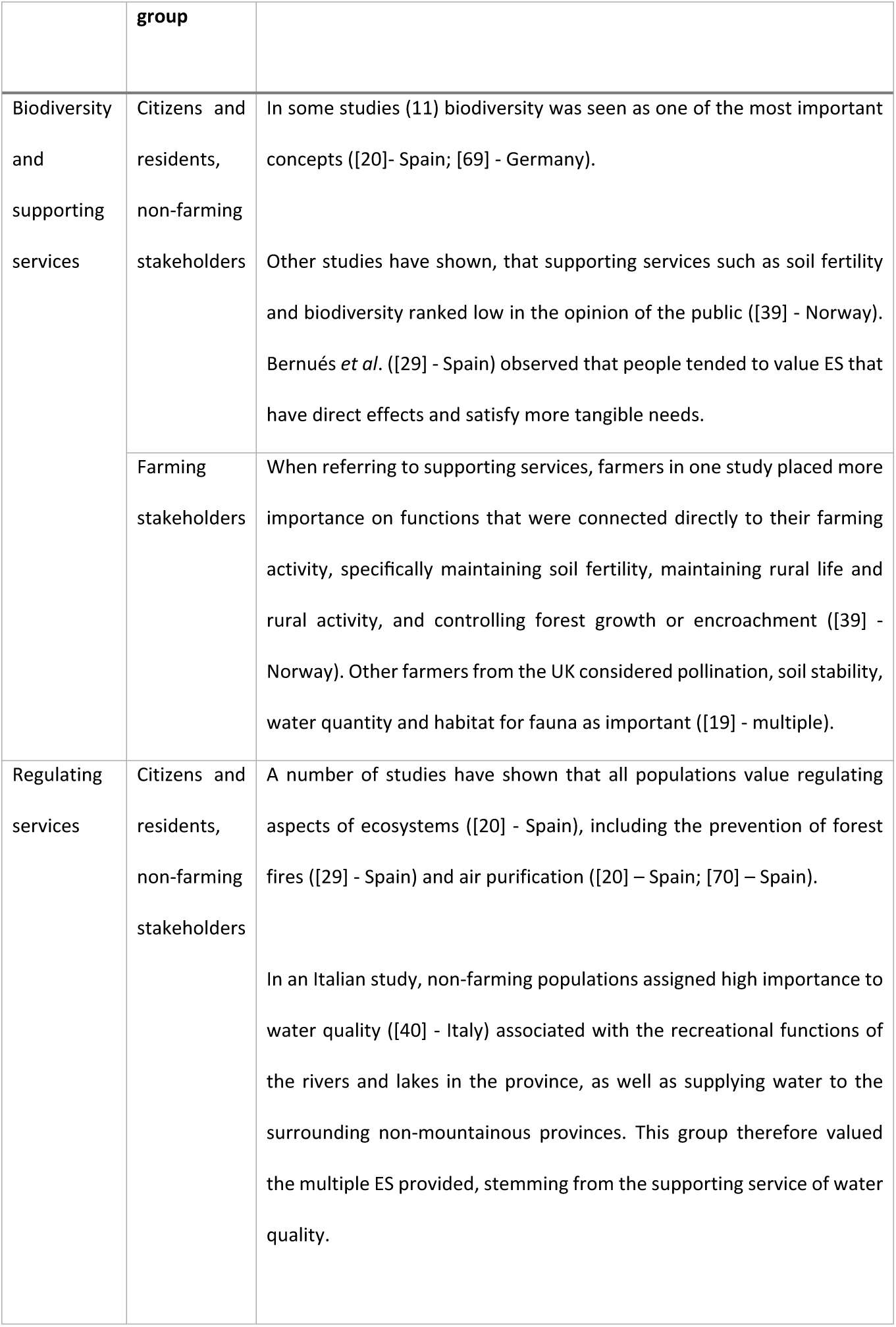
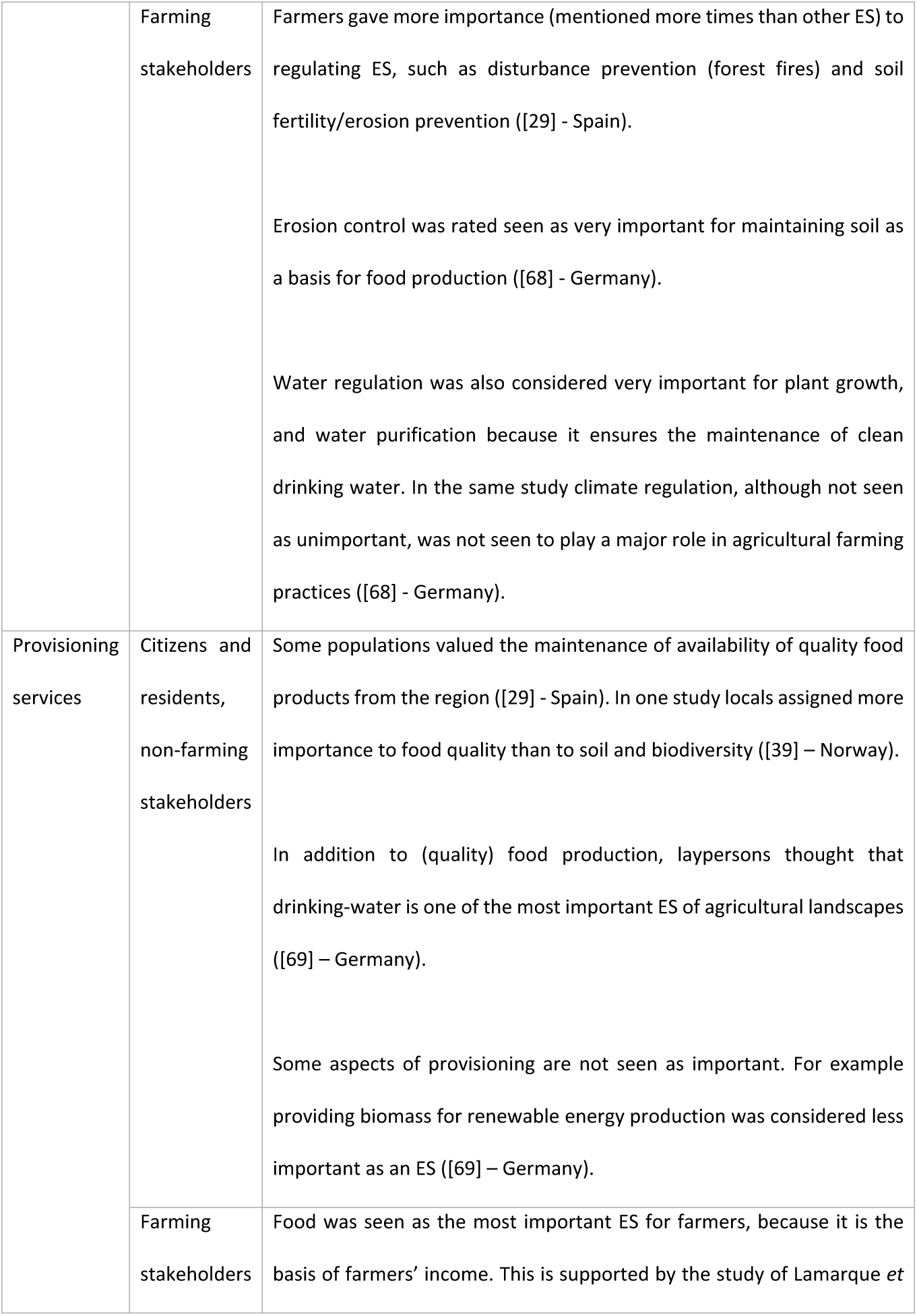
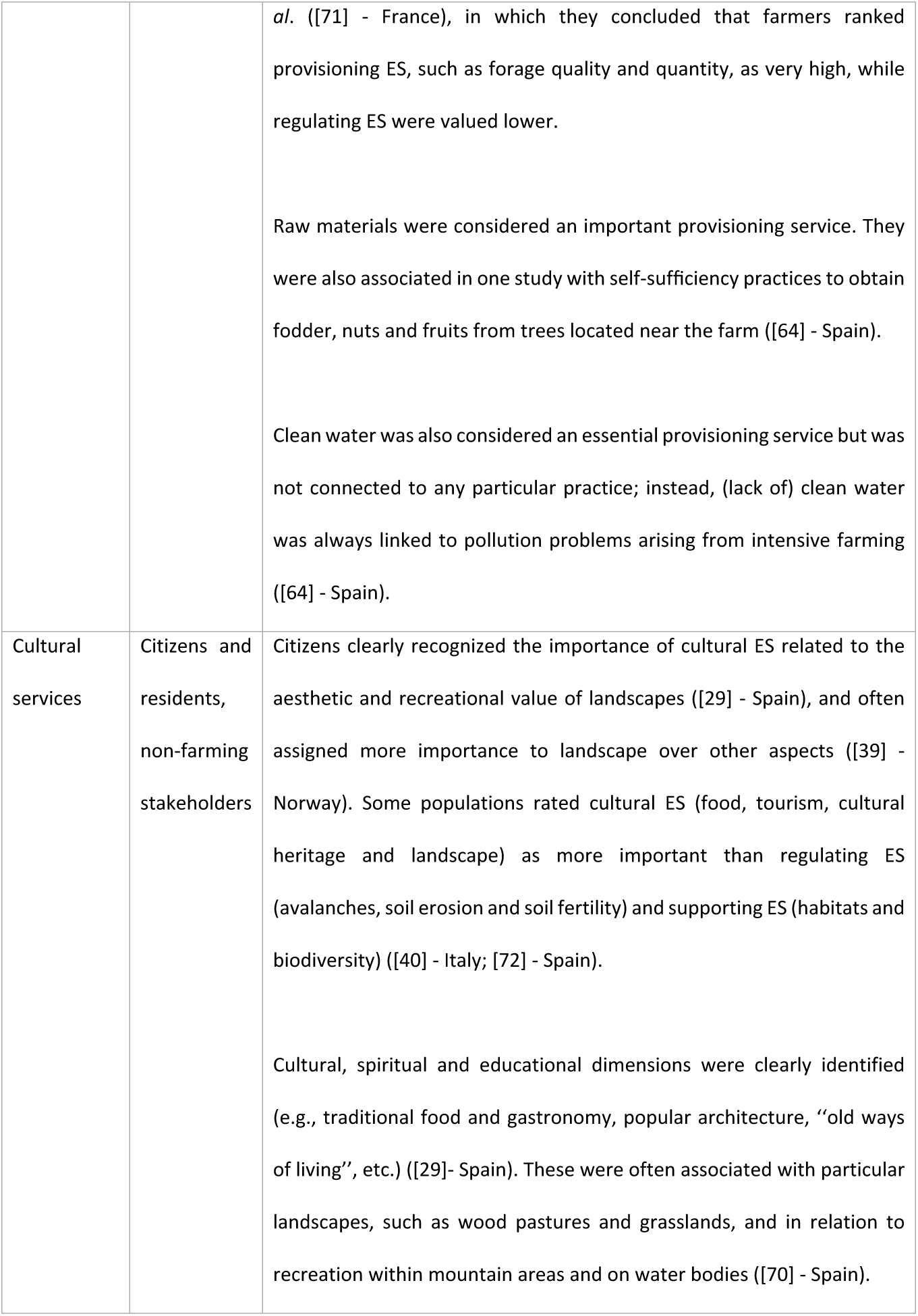
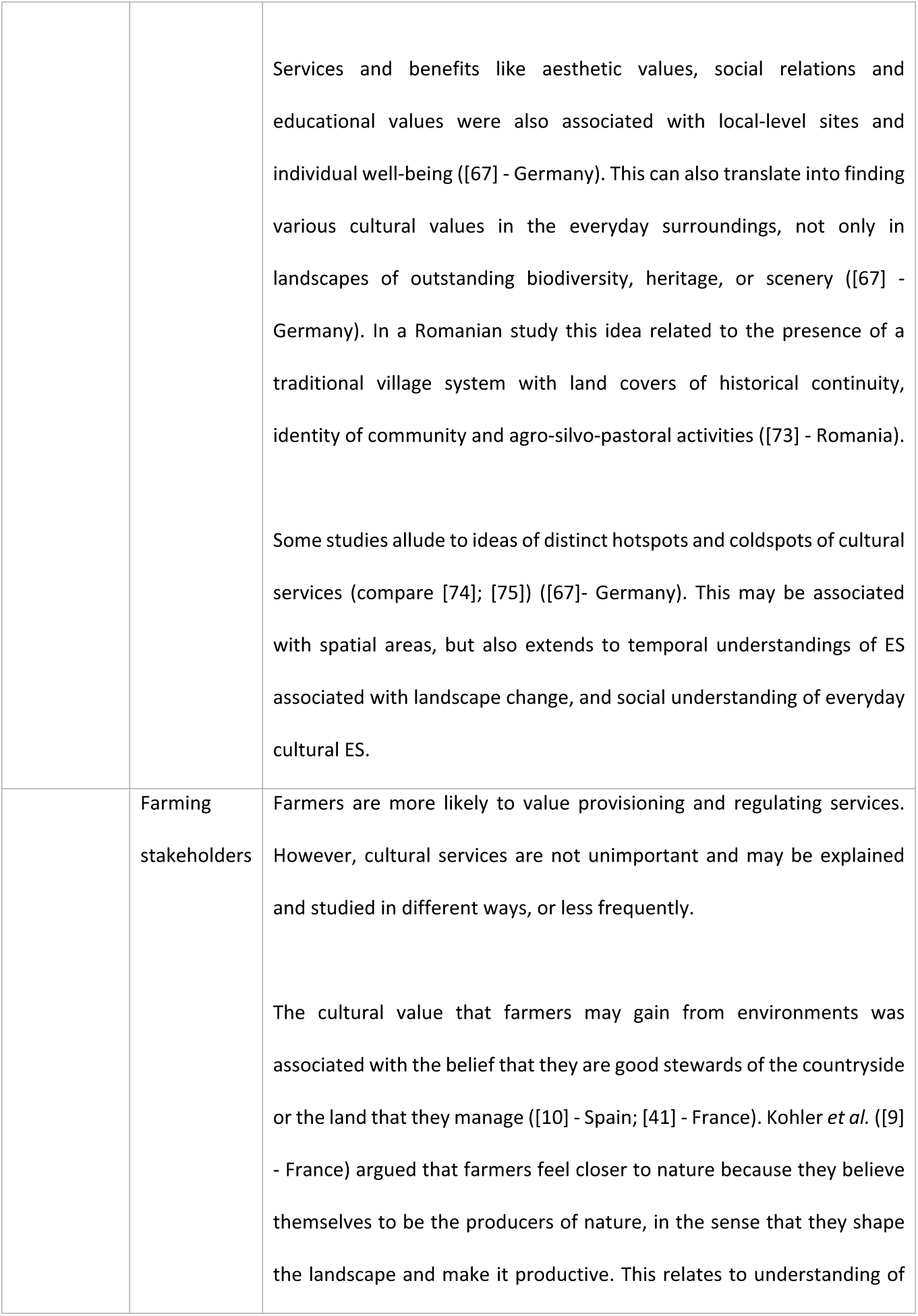
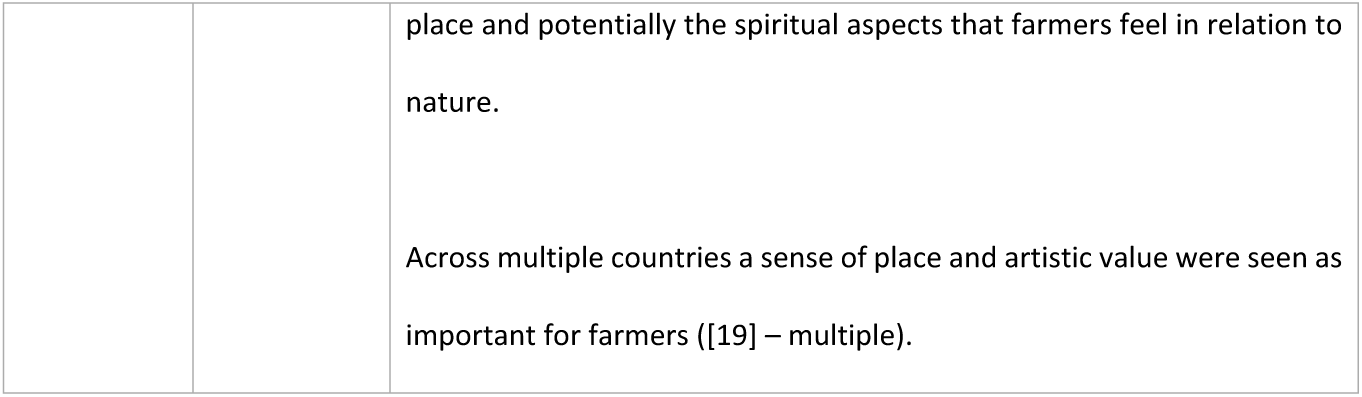
Summary of results comparing attitudes to ES from farming and non-farming stakeholder groups.

| Type of ES | Stakeholder | Summary of results |
| --- | --- | --- |

|  | group |  |
| --- | --- | --- |
| Biodiversity and supporting services | Citizens and residents, non-farming stakeholders | In some studies (11) biodiversity was seen as one of the most important concepts ([20]- Spain; [69] - Germany).<br>Other studies have shown, that supporting services such as soil fertility and biodiversity ranked low in the opinion of the public ([39] - Norway). Bernués *et al.* ([29] - Spain) observed that people tended to value ES that have direct effects and satisfy more tangible needs. |
|  | Farming stakeholders | When referring to supporting services, farmers in one study placed more importance on functions that were connected directly to their farming activity, specifically maintaining soil fertility, maintaining rural life and rural activity, and controlling forest growth or encroachment ([39] - Norway). Other farmers from the UK considered pollination, soil stability, water quantity and habitat for fauna as important ([19] - multiple). |
| Regulating services | Citizens and residents, non-farming stakeholders | A number of studies have shown that all populations value regulating aspects of ecosystems ([20] - Spain), including the prevention of forest fires ([29] - Spain) and air purification ([20] – Spain; [70] – Spain).<br>In an Italian study, non-farming populations assigned high importance to water quality ([40] - Italy) associated with the recreational functions of the rivers and lakes in the province, as well as supplying water to the surrounding non-mountainous provinces. This group therefore valued the multiple ES provided, stemming from the supporting service of water quality. |
|  | Farming stakeholders | Farmers gave more importance (mentioned more times than other ES) to regulating ES, such as disturbance prevention (forest fires) and soil fertility/erosion prevention ([29] - Spain).<br>Erosion control was rated seen as very important for maintaining soil as a basis for food production ([68] - Germany).<br>Water regulation was also considered very important for plant growth, and water purification because it ensures the maintenance of clean drinking water. In the same study climate regulation, although not seen as unimportant, was not seen to play a major role in agricultural farming practices ([68] - Germany). |
| Provisioning services | Citizens and residents, non-farming stakeholders | Some populations valued the maintenance of availability of quality food products from the region ([29] - Spain). In one study locals assigned more importance to food quality than to soil and biodiversity ([39] – Norway).<br>In addition to (quality) food production, laypersons thought that drinking-water is one of the most important ES of agricultural landscapes ([69] – Germany).<br>Some aspects of provisioning are not seen as important. For example providing biomass for renewable energy production was considered less important as an ES ([69] – Germany). |
|  | Farming stakeholders | Food was seen as the most important ES for farmers, because it is the basis of farmers' income. This is supported by the study of Lamarque *et* |
|  |  | *al.* ([71] - France), in which they concluded that farmers ranked provisioning ES, such as forage quality and quantity, as very high, while regulating ES were valued lower.<br>Raw materials were considered an important provisioning service. They were also associated in one study with self-sufficiency practices to obtain fodder, nuts and fruits from trees located near the farm ([64] - Spain).<br>Clean water was also considered an essential provisioning service but was not connected to any particular practice; instead, (lack of) clean water was always linked to pollution problems arising from intensive farming ([64] - Spain). |
| Cultural services | Citizens and residents, non-farming stakeholders | Citizens clearly recognized the importance of cultural ES related to the aesthetic and recreational value of landscapes ([29] - Spain), and often assigned more importance to landscape over other aspects ([39] - Norway). Some populations rated cultural ES (food, tourism, cultural heritage and landscape) as more important than regulating ES (avalanches, soil erosion and soil fertility) and supporting ES (habitats and biodiversity) ([40] - Italy; [72] - Spain).<br>Cultural, spiritual and educational dimensions were clearly identified (e.g., traditional food and gastronomy, popular architecture, “old ways of living”, etc.) ([29]- Spain). These were often associated with particular landscapes, such as wood pastures and grasslands, and in relation to recreation within mountain areas and on water bodies ([70] - Spain). |
|  |  | Services and benefits like aesthetic values, social relations and educational values were also associated with local-level sites and individual well-being ([67] - Germany). This can also translate into finding various cultural values in the everyday surroundings, not only in landscapes of outstanding biodiversity, heritage, or scenery ([67] - Germany). In a Romanian study this idea related to the presence of a traditional village system with land covers of historical continuity, identity of community and agro-silvo-pastoral activities ([73] - Romania).<br>Some studies allude to ideas of distinct hotspots and coldspots of cultural services (compare [74]; [75]) ([67]- Germany). This may be associated with spatial areas, but also extends to temporal understandings of ES associated with landscape change, and social understanding of everyday cultural ES. |
|  | Farming stakeholders | Farmers are more likely to value provisioning and regulating services. However, cultural services are not unimportant and may be explained and studied in different ways, or less frequently.<br>The cultural value that farmers may gain from environments was associated with the belief that they are good stewards of the countryside or the land that they manage ([10] - Spain; [41] - France). Kohler *et al.* ([9] - France) argued that farmers feel closer to nature because they believe themselves to be the producers of nature, in the sense that they shape the landscape and make it productive. This relates to understanding of |
|  |  | place and potentially the spiritual aspects that farmers feel in relation to nature.<br>Across multiple countries a sense of place and artistic value were seen as important for farmers ([19] – multiple). |

#### 3.3.1. Conflicts and trade offs

Often conflicts can appear between different perceptions of ES. However, few studies covered conflicts of interest, as most focused on understanding certain populations’ views. Conflicts were found to arise between multiple groups ([76] – Hungary). For example in Hungary, during the development of a national park, trade-offs were made between cultural services and decreased provisioning services. Kovács *et al*. ([76] – Hungary) state that the national park service, the local inhabitants, visitors, scientists and students who pursue cultural, supporting and regulating services were the winners of the changing situation, while the losers were the local farmers who had a more production-oriented opinion, and who value ES such as food from agriculture over regulating or cultural services.

Lamarque *et al*. ([71] – France) found that visions of ES differed between farmers and scientists. Farmers in the study area explained that, for them, ES are not numbers or aspects to be measured, but part of a complex system of decision-making. Pietrzyk-Kaszyńska *et al*. ([77] – Poland) reported that conflict can be internal to a stakeholders group. For example, the opinions of landowners or farmers can be contradictory in relation to protected areas; some can declare their love for nature and at the same time support activities that cause its degradation [78].

### 3.4. Attitudes to landscape and landscape change

Attitudes to landscape and landscape change were an important focus in the papers studied. Seventeen papers focused on the aesthetic quality of the landscape, often asking non-farmer populations about preferences for certain landscape types, sometimes using visual cues, scenarios and photographs to understand choice and preference, sometimes using field trips or field studies to gauge opinion. Many studies focused on whole landscape scale and often included grassland as part of the landscape. Some studies aimed to better understand farmers’ views about aesthetics of agricultural landscapes. However, there is an immense variability in visual landscape preferences among groups ([79]– Spain). This also includes variability within groups and between regions and contexts.

In relation to whole landscape aesthetics, Schirpke ([80]– Austria) found that amongst non-farming populations, high aesthetic value was given to terrestrial viewpoints mainly located at high altitude, allowing long vistas, and including views of lakes or glaciers, and the lowest values were for viewpoints close to streets and in narrow valleys. Water related features were often seen as a dominant attribute in terms of visual amenity ([81]– Ireland). In relation to a specifically agricultural context, Bernués *et al*. ([64]– Spain) showed that aesthetics was seen to be one of the most important ES. According to the non-farmers, aesthetic value was affected by avoiding the overexploitation of pastures (to prevent desertification); grazing in the mountains (to create and maintain meadows); maintaining traditional buildings (to provide shelters, water points, and popular architecture); and freely grazing animals (to enhance the quality of the landscape, to make it more beautiful), and to offer spiritual experience to those who value contact with animals (see also [14] – Switzerland).

Some studies focused on understanding the characteristics of highly valued landscapes. Häfner ([60] – Germany) found that about 70% of (non-farmer) respondents preferred diverse and highly structured landscapes, which is a common finding in other studies, and included a clear preference for crop diversity, point elements and linear elements. Point elements, such as a single tree within the fields out-value other attributes. Linear elements are seen to be valued, such as high-stem fruit trees, hedgerows and low-intensity pastures with trees and bushes ([14]-Switzerland). López-Santiago ([72] – Spain) also showed that the linear element of a ‘drove road’ (traditional path through the landscape for transhumance populations to move livestock) was highly valued, partly due to its cultural associations, but also its aesthetic value. A third of the population in Häfner’s study seemed to prefer ‘cleared-out’ landscapes with very low degrees of complexity. Although Häfner’s study does not explore why people hold these views, it does explore influential factors, for example ‘political opinion, ‘preferred outdoor activity’ and ‘means of transport’. Howley ([81, 82]– Ireland) reported that their respondents preferred a mixed farming landscape, and extensive farming landscapes over the more modern intensive farming landscapes (also see [83] – UK) due mainly to the homogeneity of more intensive landscapes [84]. Landscape preferences were also affected by factors such as seasonal stage (e.g. flowering stages are valued highest). Equally, naturalness and diversity were important to non-farming populations ([14] – Switzerland), particularly in terms of species richness in grassland ([62] – Germany).

Farmers can sometimes hold opposing views to non-farming populations. For example, Junge *et al*. ([85] – Switzerland) found that whilst non-farming populations preferred a mixed landscape with 30% of the land used for ecological compensation areas (ECAs), farmers preferred cropland with 10% ECAs. This study not only demonstrated the differences between the populations in terms of attitude, but also that the agri-environment schemes affected the visual attractiveness of the landscape through a change in the land use patterns. Andersson *et al*. ([86] – Sweden) found that there were differences in expectation between farmers around visual preferences following different intensity regimes. For example, high-intensity farmers saw the importance of having well-tended farms, where the land, buildings and infrastructure should be in good condition. They were interested in the land looking, and being, economically valuable. Equally, Junge *et al.* ([85] – Switzerland) and Kohler *et al*. ([9] – France) found that some farmers showed a strong preference for neat, clean and ordered landscapes. Junge *et al*. ([85] – Switzerland) report that this might have been due to these farmers having a strong internalized sense of ‘stewardship and care’ and/or believing that production was the most important function of agriculture, over and above ecological function. Such views of aesthetics can cause conflict when conservation often creates environments that may be viewed as untidy or messy (Kovács *et al*., [76] – Hungary). Tidiness, however can also be perceived as positive by non-farmers, for example regional experts in an alpine environment perceived a tidy landscape to be aesthetically pleasing, partly due to associations with avalanche regulation ([19] – multiple).

Quétier, *et al*. ([41] – France) suggest that although aesthetics is an important element of landscape management it often favours “postcard” or heritage social representations. In addition Willis ([83] – UK) showed that people prefer the status quo landscape, which has consequences for any need or drive for landscape change. Konkoly-Gyuró ([87] – France and Germany) highlighted that landscape can be seen in very different ways: as a photo, a snapshot of reality, and/or as a film, or a dynamic entity. As such it has links to the past and the future, to images of stability and change, where conflict and degradation are associated with change. In an earlier study, Marcel *et al*. ([88] – Switzerland) suggest that it is not landscape change that is assessed as good or bad, but the (related) change in the meaning of the landscape elements that lead to positive or negative assessments. In an Italian study, major driving forces of landscape change were perceived to be planning policies, technological innovation, agricultural policies and demographic trends ([89]– Italy). In a Romanian study, Pătru-Stupariu *et al.* ([73] – Romania) found that causes of landscape change were: increasing tourism, land tenure and social changes, land-use intensification, post-communist transition, and foreign investments. In Spanish savannah grasslands (dehesas), concerns related to a lack of regeneration, oak decline (oaks are part of the traditional diverse land use of dehesas), and conversion to urban areas ([10] – Spain).

### 3.5. Attitudes towards agricultural management practices

#### 3.5.1. Impact of management practices

Attitudes to particular agricultural management practices can vary within stakeholder groups as well as between farming and non-farming populations. For example, conservation practices are viewed differently by different groups of farmers. Fish *et al*. ([90] – England) show that for some farmers the idea of conservation was thought to be incompatible with what they regarded as the dynamic nature of landscapes, “freezing” landscapes at particular moments in time when they should be allowed to modify and change incrementally. In relation to particular management practices for grassland, the utility of practices such as sowing more diverse seed mixtures can be perceived differently by farmers and conservationists/researchers. For example, a proposal by researchers to sow more diverse seed mixtures in France to improve biodiversity ([91] – France) did not interest the farmers. This was because their primary goal for the grassland was hay-making and they saw no advantage in increasing the number of species beyond four for this purpose. Similarly seed mixes designed to improve pollination ([92] – England) were seen as effective by farmers, but also costly because of the lengthy preparation process.

Perceptions of cost can also manifest in lack of uptake of other management practices for grassland. Collins *et al*. ([93] – England) report the results of a survey looking into measures most likely to be implemented in the future by farmers. The list of measures included many low cost or cost-neutral measures, suggesting that costs continue to represent a principal selection criterion for many farmers. Other perceptions of impact of management measures can have an effect on management uptake. For example, the adoption of non-inversion tillage across four European countries was seen as lower for dairy farmers because it was perceived to result in lower yields, more weeds, more crop diseases, and to affect the way the fields would look ([94] – Belgium, the Netherlands, Germany, Italy). One positive driver of uptake however was increased soil carbon, although this effect is often small and inconsistent. Feliciano *et al*. ([95]– Scotland) found that grassland farmers in particular are proud of the role that grassland management plays in storing carbon and believe they should be more clearly recognised for their role in climate change mitigation.

#### 3.5.2. Policy, regulations and agri-environment schemes (AES)

Attitudes to policies relevant for land management were acknowledged as a factor affecting land management decision-making in many studies. Perceptions of policies and AES were often studied in relation to uptake. Some studies demonstrated the often negative perceptions farmers can have of environmental policies. For example, Alavoine-Mornas *et al*. ([96] – France – Alps) showed that whilst most farmers pay attention to nature conservation, they can often perceive nature conservation policies as restrictions, bans and limitations. A similar finding was reported from high intensity farms in Sweden, where farmers often viewed regulations on preservation of cultural heritage as irritating (Andersson *et al*., 2015 – Sweden). Such policies may not have been taken up by high output farmers as they may have felt the policies threatened their independence and took away the pride they have in producing crops from their land. Fish *et al*. ([90]– England) described the motivation of grassland managers to participate in AES as tied up in their economic perception of land, the importance of profitability, and their economic concerns as a mechanism for achieving conservation goals.

Bernués *et al*. ([64] – Spain) demonstrated the value of AES to livestock farmers. For example, sheep farmers perceived that they can maintain profitability through participation in AES. Beef farmers perceived that subsidies allowed them to remain a viable business in the face of meat price changes. Such farmers were also concerned about the way that subsidies were designed including the horizontal distribution of premiums that does not account for differentiation according to the character of the farm ([64]– Spain; Schroeder *et al*. 2015 – England). Such concerns were also manifested in grassland farmers participation in AES for nature conservation in Germany ([97] – Germany), where only a minority of grassland farmers were using the existing AES to the full extent. This was attributed to a perception by 40% of the farmers that AES are not an important tool to preserve the landscape and nature. This is similar to previous studies such as Fish *et al*. ([90] – England), where some land managers believed that AES schemes did not make any enduring mark on the landscape. Despite the majority of grassland farmers having a positive attitude towards nature conservation, this does not automatically lead to a positive attitude towards participation in AES ([97] – Germany).

Non-farming stakeholders also have perceptions of AES. For example, institutional and civil society stakeholders in Italy acknowledged the importance of EU policies supporting farmers’ income and activities in rural areas in driving impact on landscapes ([89] – Italy). However, both these stakeholder groups also raised doubts about the real effectiveness of AES in terms of containing farm land abandonment in marginal areas.

#### 3.5.3. Innovation and technology

Some studies of management practices included the adoption of new technology. Such studies have found that attitudes towards a technology have a strong influence on whether or not farmers intend to adopt new technologies ([98] – England). For example, perceptions held by dairy farmers around the usefulness and ease-of-use of a technology were likely to influence their adoption of technologies relating to grazing, artificial insemination and financial management, particularly when they are primarily motivated by financial considerations ([99] – EU and Ireland).

Some studies found that the attitudes and perceptions about farming in general also influenced local residents and the general population’s attitude to technology, particularly in connection with food products produced from managed farmland, including grassland. For example, Rodríguez-Ortega *et al*. ([100] – Spain) has shown that people with a productivist profile (agree with maximising profits from natural resources, using intensive agriculture to solve hunger, maintaining agricultural policies and premiums based on production), were more likely to agree that new food technologies played a reassuring role in improving food quality and safety. This is in comparison to those with a conservationist profiles (opposite to productivist) who were less likely to agree that technology improved product quality and food safety. The opinions of the consumers of products is therefore also valuable when considering grassland management and landscapes.

### 3.6. Factors affecting attitudes

#### 3.6.1. Contextual influences

A number of factors were investigated as potentially influential or partly explanatory of the attitudes and preferences of different stakeholders. These included age, income and other economic factors, biophysical factors in the landscape, socio-cultural background, farm structure, education, and gender, (Table 4). These factors were mentioned in relation to the influence on behaviour or intentions.

**Table 4.** Summary of contextual themes affecting attitudes to landscapes.

| Theme | Example findings |
| --- | --- |
| Age | Older people tended to prefer intensive and mixed farming landscapes, as well as traditional farming landscapes perhaps due to familiarity, and younger people had a stronger preference for water related landscapes ([81]– Ireland; [82]- Ireland)<br>Older people were more likely to perceive important services to include local ecological knowledge, nature recreation activities, soil fertility and erosion control, fire prevention, pollination, and gathering of crops. Younger people more often |
|  | perceived food from agriculture, air purification, habitat for species and the genetic pool as important services. This was seen as a combination of changing needs throughout life and a changing engagement with knowledge about ecosystems through the educational system ([101] – Spain).<br>Younger people in the general population and local residents, as well as younger farmers ([102] - Belgium), were seen to have more positive attitudes to nature conservation ([100] - Spain).<br>Age was seen to influence the level of land manager satisfaction with designation of protected sites under Natura 2000. The oldest and youngest groups were the most satisfied, with those aged 40-59 the least satisfied ([77] – Poland).<br>Age also affected land manager appreciation of land uses such as oak ([10] - Spain); and relative preferences for wild vs managed nature scenes ([103] – Netherland). |
| **Income and other economic factors** | Fixed monthly income of farmers is a factor influencing their allocation of income to public goods ([104]– Italy).<br>The share of farm income in total household income plays a negative role on farmers' marginal participation in all AES ([105] - Italy).<br>Farmers with low adjustment abilities find farming is a more rewarding job in terms of quality of life, independence, lifestyle, than it is in terms of money ([106] – multiple). |
|  | Economic incentives might influence a ‘productivist’ farmer to take the climate into consideration ([107] – Ireland).<br>The failure of high input agriculture to lead to financially viable farms has led to more positive beliefs about decreased input use, and conversion to organic farming; ([108] – Scotland).<br>A high proportion of rented farmland could have a negative influence on the farmers' perception of semi-natural features and on their willingness and commitment into schemes aimed at biodiversity preservation. These farmers might be reluctant to engage in environmental schemes on agricultural parcels which could not be part of their farmland in a few years ([96]:37 - France). |
| **Biophysical factors in the landscape** | Conservationists associated high soil fertility with high conservation value of land, and favoured the protection of birds. Farmers associated a high conservation value with vegetation units displaying high plant species richness ([18]– Germany).<br>Physical characteristics of landscape elements are seen as stronger predictors of aesthetic preference than sociodemographic variables of study participants ([14] – Germany).<br>The state of grasslands (abandoned/well managed) or landscape diversity and fragmentation (presence of hedges, trees or a stream) were sometimes discussed by stakeholders as important elements linked to ES such as habitat for fauna or aesthetic value ([19]– multiple). |
| **Socio-cultural background** | The perception of ES has shown to vary according to socio-demographic factors and individuals' backgrounds and interests. For example, stakeholders directly involved in agriculture or the products or services of livestock farming, perceive the ES more positively. In an Alpine context, local populations of farmers did not perceive the livestock sector to be a threat for water quality and climate change, however non-farmers were uncertain ([40] – Italy).<br>Respondents in a relatively higher social classes (UK National Readership Survey social grades) were found to be less likely to rate mixed farm landscapes highly and more likely to rate wild nature scenes highly than respondents in relatively lower social class groupings ([81] - Ireland).<br>Farmers, older respondents, and respondents with low levels of income and education displayed relatively low preferences for wild natural landscapes, while respondents with a preference for green political parties, younger respondents, and respondents with high levels of income and education, displayed relatively high preferences for wild nature ([103] – Netherlands).<br>Regional variables are seen to have an influence on preferences for various landscape categories, with residents' preferences for mixed farming landscapes and water related landscapes explained significantly by their 'rural' location, perhaps due to familiarity ([81]– Ireland).<br>Appreciation of ES and transhumance landscape preferences were seen to vary within four stakeholder groups, including people rooted in transhumance, people who were |
|  | environmentally aware, urban inhabitants with recreational needs, and farmers with strong cultural identity ([72] – Spain). |
| **Farm structure** | Farmers with a more extensive grassland management (low nitrogen input, lower stocking rates, and low cutting frequency) showed a more positive attitude towards AES. Farmers with larger farms were more likely to take part in AES due to the fact that compensation payments are paid per ha and take into account marginal land ([97] – Germany)<br>Farm structure and size interacts with farmer perceptions and attitudes. For example, in combination with other factors such as higher number of years worked in farming, membership of farmers’ unions, and less time in formal education, farm size can link to indifference to the policy context e.g. subsidies and income support ([106] - Slovakia and Lithuania)<br>Findings indicated higher climate change awareness with increasing farm size, although farm size was also associated with region, and attitudes may have been influenced by the climatic conditions (e.g. sub-continental and drier summers) ([109] – Germany) |
| **Education** | Education is often associated with other factors to influence preference. For example, an increased (positive) sensitivity was seen within particular groups for an element-rich and diverse landscape. These included women, university graduates and people with a pronounced environmental value setting ([60] – Germany)<br>Both higher education and membership in an environmental organisation were |
|  | predictors of a dislike for landscapes without ecological compensation areas (ECAs), probably due to more knowledge about the ecological benefits of agro-biodiversity ([85] – Switzerland). |
| **Gender** | Gender differences emerge in relation to ES. For example, men tended to consider the most important ES to be those related to raising livestock (i.e., livestock, manure, seed dispersal and bullfighting events), and women typically perceived regulating services (i.e., tree regeneration and soil erosion control) as more valuable. These have been explained in relation to gender-differentiated environmental awareness and the gender division of work ([101]– Spain).<br>Individuals who are older and/or female were found to have a stronger preference for traditional farm landscapes ([82] – Ireland).<br>Gender difference in attitudes towards nature conservation were reflected in the fact that conservation had greater support among females (in the general population) ([100] – Spain). |

Contextual factors can be both intrinsic (gender, age, education) and extrinsic (farm structure, income, biophysical landscape). Each combine to create a context from which individuals and collective groups of stakeholders form attitudes. Table 4 indicates that some factors have been shown in studies to affect both farming and non-farming stakeholders, including age, biophysical landscape and socio-cultural background. Other factors have been mostly associated with either farmers (farm structure and income), or non-farming populations (gender and education). It is worth noting that the examples shown in Table 4 only include instances where notable or statistically significant links between the themes and attitudes have been found. These contexts are highly variable, as are the circumstances in which links to such themes are not significant.

#### 3.6.2. Knowledge (tacit, learned, experiential, local)

Stakeholder perceptions can be associated with multiple forms of knowledge. For example, Leroy *et al*. ([61] – Multiple) showed that perceptions are influenced by local knowledge (localized, experiential or indigenous knowledge), scientific knowledge (explicit knowledge that has been derived from applying more formal methods; Raymond *et al*., 2010), as well as the level of involvement in management of local environment and resources ([19] – multiple; [61]– multiple).

Oteros-Rozas *et al*. ([101] – Spain) identified local ecological knowledge as a key variable for explaining the different perceptions amongst stakeholders of the important ES, including prevention of soil erosion, that are delivered by transhumance livestock systems (moving animals between summer and winter pastures). Bernués *et al*. ([39] – Spain) also demonstrated that in an assessment of multi-functionality of fjord and mountain agriculture, soil fertility was important for farmers, who were more aware of the central role of soil fertility in agricultural production and the delivery of other ES, compared to city dwellers. Sattler and Nagal ([38] – Germany) also agreed that experiential knowledge and acquired knowledge affect attitudes, which manifest as risk tolerance, environmental awareness, innovativeness and adaptability in individuals.

A lack of knowledge can be restrictive, e.g. a lack of technical knowledge and experience was seen to negatively affect the adoption of non-inversion tillage by Belgian dairy farmers ([94] – Belgium, the Netherlands, Germany, Italy). Equally, a lack of evidence (for example linking specific farming practises to water or air quality outcomes) can affect farmers’ attitudes and therefore their behaviour ([93] – England). Information can also lead to transformative processes such as land conversion in national parks. For example, Hjalager *et al*. ([110] – Denmark) showed that the more that is known about the goals and organisational aspects of national park projects, including new business opportunities, the more confident farmers with protected land in the national park can be about converting land to more sustainable practices.

Sources of knowledge are variable and influential in different ways. For example, Dietze *et al*. ([68] – Germany) showed that important sources of knowledge and information for farmers forming opinions and decisions about soil ES in agricultural management are education and training through university or extension services. They found that no farmers received information through exchanges with specialists or academic journals, despite other studies finding that scientific knowledge can be needed by farmers to legitimise their own knowledge ([91] – France). Significantly, Dietze *et al*. ([68] – Germany) found that the most important form of knowledge for decision-making around the adoption of new agricultural practices was long term experience, both individual and passed between generations, as well as among colleagues and neighbours, and producers of agricultural machinery. In a study of green payments and perceived rural landscape quality in Italy, Cortignani *et al*. ([104] – Italy) demonstrated that knowledge of environmental and landscape policies was also a factor relevant for increasing landscape quality maintenance.

Particular management interventions can affect opinions of the landscape between stakeholders. For example, in a study of landscape change in Switzerland, Hunziker *et al*. [88] identified that experts and lay people had a different opinion of landscape. The experts rated a traditional cultural landscape much higher than lay people, who rated reforestation and intensification much higher than the experts. The experts’ opinion was likely to have been influenced by the long-lasting effect of a program explicitly promoting maintenance of the traditional cultural landscape in the Alps. There can also be differences between local farmers and experts linked to knowledge. For example, Lamarque *et al*., ([19] – France, Austria and UK) showed that differences between ES considered important by regional experts and local farmers within regions appeared to reflect differences in technical and local knowledge (generated by practice and observations). These could be reflected in different objectives and concerns leading to divergent priorities among stakeholders for ecosystem management. Lamarque *et al*., ([19]– France, Austria and UK) concluded that increased awareness of the utility of particular services for sustainable management might be needed [111].

#### 3.6.3. Environmental and personal values

In a study of public preferences for traditional farming landscapes, Howley *et al*. ([82] – Ireland) showed that personal characteristics and environmental value orientations strongly influenced preferences for all the landscape types. Particular environmental value orientations were found to be influential, including ‘ecocentric’, where individuals are likely to be against even a very small degree of human intervention on landscapes; ‘anthropocentric’, where people hold a human-centred value for the environment [112], ‘agricultural productivist’, in which the capacity of the landscape for producing food and fibre make mixed farming landscapes more attractive than wild nature scenes to some. Negative environmental value orientations also continue to be significant, including apathy, which has been shown to be negatively associated with a preference for wildlands and for cultural landscapes [113]. Such values may be expressed in political opinion, where, for example, preference for green political parties can relate to preferences for wild versus managed natural areas ([103] – Netherlands).

Experiences often combine with values to influence attitudes. For example, López-Santiago ([72] – Spain) showed that cultural attachments or previous experience were important for differences in the visual perception of ES supply, and preference for transhumance landscapes. Equally, experiences of outdoor activities and transport ([60] – Germany), and experiences of care and stewardship ([114] – Italy) were shown to be influential on landscape and land use preference.

Personal goals and family values can also be important for both farmers and non-farmers. For farmers, the drivers for particular preferences may be associated with personal development and family life goals beyond the farm enterprise ([115] – UK), or the existence of children or family members employed on the farm ([38] – Germany). Equally, for land managers, preferences for particular landscapes might be driven by the potential for the land to provide real estate and continue family tradition, for example the preference for oak conservation in the dehesa grasslands in Spain ([10] – Spain). Other non-farming stakeholders have been shown in previous studies to be influenced by family values. For example, in rural landscape planning in Italy, civil society stakeholders’ opinions were formed based on the needs of their own family, and they often could not easily assess the landscape scale [89].

### 3.7. Differences between farmer and non-farmer attitudes

Table 5 summarises the attitudes to grassland, landscapes, agricultural management and ES delivery on both non-farming (citizens, consumers, residents, locals, public, visitors, experts) and farming (farmers and land managers) populations studied in the papers reviewed. It shows that attitudes can vary between stakeholder groups based on the experience, background, context and characteristics of each stakeholder group. Such contrasts highlight potential trade-offs that need to be considered when making decisions about land management, particularly relevant to environments containing grassland landscapes.

**Table 5.** Summary of attitudes, and influences on attitudes, for non-farming and farming stakeholders reported in the papers included in the systematic review.

| Attitudes | Non-farming stakeholders | Farming stakeholders |
| --- | --- | --- |
| Attitudes to grassland/<br>grassland<br>management | Public have a knowledge of, and interest in, various benefits provided by specialist grasslands; they have aesthetic appreciation of species-rich grassland; varied opinions around presence or lack of grazing livestock; local populations can | Farmers, may have complex attitudes towards grasslands and management based on the complexity of trade-offs; appreciate benefits such as forest fire prevention; differences in attitudes to grazing; can prefer intensively used grasslands and value soil fertility and |
|  | value grassland as a resource, compared to visitors who value aesthetics more strongly. | conservation over carbon sequestration; increased grassland can be seen as an administrative burden. |
| Attitudes to ecosystem services | Benefits of cultural ES, including opportunities for recreation, spiritual and cultural experiences and the provision of food, mainly relating to quality and safety issues; sometimes biodiversity and soil fertility are important, but sometimes less important due to lack of perception of direct benefits; high importance for regulating services when they help to maintain recreation or a beautiful environment; experiences and tradition were important. | Placed more importance on ES that were connected directly to their farming activity; ES such as soil fertility, water quality and productivity are important for maintaining a viable farm system; production of food or forage is highly rated as it relates to the farmers' income and livelihood; less likely to comment on cultural services, but may be understudied. |
| Attitudes to landscape and landscape change | Value long vistas and elements such as grazing animals, water and traditional buildings; prefer diverse and highly structured landscapes – often with low degrees of complexity; mixed farming and extensive farming with more naturalness; prefer the status quo. | Some farmers prefer the presence of some ecological 'natural' areas, but fewer than non-farmers; high-intensity farmers prefer well-tended farms, where the land, buildings and infrastructure look economically viable; some farmers prefer neat, clean, ordered landscapes, reflecting stewardship. |
| Attitudes towards management practices | Expert stakeholders perceive policies as key for supporting farmers, rural activities and impact on the landscape; few studies assess the opinions of public on management practices or policies; in relation to technology those with | Attitudes to management options reflect each farmers perceptions of the core purpose of farming; cost is an important influence; perceptions of outcomes is important; some farmers see nature conservation policies as restrictions, bans and limitations rather than |
|  | conservationist views are less likely to perceive positive impacts of agricultural technology on food quality and safety than those with productivist views. | opportunities; some beef farmers believe subsidies allow them to stay viable; perceptions of utility and ease of use are important for technology uptake. |
| Factors affecting attitudes | Age was a factors affecting aesthetic appreciation and perceptions of conservation; conservationists were influenced by biodiversity; physical state of grassland affects attitudes to ES; social class contributes to perception of aesthetic appreciation of mixed farms and wild nature scenes; rural dwellers prefer mixed farming landscape; women and university graduates prefer diverse and traditional farm landscapes; men prefer ES related to raising livestock; women prefer ES related to regulating services; knowledge of policy impact, and technical and local knowledge can affect expert attitudes to important ES; environmental value orientations, experiences and family values influence preference for landscapes. | Age affected perceptions of conservation; income and economic costs affect attitudes to management, climate change and view of farming; farming background affects attitude to livestock farming, environmental impacts and links to lower preference for wild natural landscapes; farm size and types links to participation in AES and attitudes to climate change; knowledge of ecosystem and farming processes shape attitudes to functionality and management of land; lack of knowledge can lead to lack of uptake of management; experiential knowledge is highly valued; technical and local knowledge can affect expert attitudes to important ES; personal and family goals affect attitudes to landscapes and management. |

## 4. Discussion

This literature review summarises the breadth of literature on attitudes and perceptions to grassland landscapes and associated ES and management practices. It shows that relatively few studies focused directly on grassland or PG. Where attitudes were a focus, studies tended to be around specific grassland types or systems such as meadows, species-rich environments, and alpine grassland areas. Because context was important in the majority of studies reviewed, it would be useful for future studies to focus more directly on attitudes to grassland in a greater variety of contexts. Equally, grassland is not always encompassed within an individual issue and cannot be easily separated from other agro-ecological contexts. Attitudes and perceptions vary greatly between and within stakeholder groups due to the variety of environments and the experience of the stakeholders. Each grassland environment is used or perceived uniquely by each group of stakeholders. Farmers in particular, may have complex attitudes towards grasslands and management often based on the complexity of trade-offs they have to make between profitability and ES delivery. It is likely, however, that non-farming stakeholders also hold complex attitudes. They are able to focus on the benefits of grassland environments and perceive positive impacts of particular management regimes or land use patterns, albeit in different capacities across and between groups.

The majority of studies in the review focused on Western European farming environments. Countries missing in representation were those at the extremes of the continent, such as Latvia, and Portugal. Future research in the less studied farming environments in Eastern Europe would complement and expand on the studies clustered in Central and Western Europe. Most of the expected populations and interest groups were included in the review, but what might be missing from the map of influential factors is a better understanding of the view and actions of consumers, and the influence of markets and supply chains (which was not a distinct target in search criteria for this review). However, some of this information has been found and reviewed in a corresponding systematic review of economic drivers of farmer adoption [116]. A link to the understanding of consumer attitudes as a driver for farmer decision-making could be investigated in future studies.

Limitations of the systematic review include the exclusive focus on English language, which may have limited the accessibility of some studies. For example, specific considerations on *dehesa* and *montado* grassland in the Mediterranean region may have been omitted (e.g. [117-119]). Other limitations include the omission of the date of data collection from 32 papers. However, despite limitations, the study as a whole has begun to identify the context in which PG is managed across Europe and is representative of the type and balance of studies that exist, the study also helps in identifying where more targeted searching could reveal in depth information and research on attitudes to PG landscapes and management.

In order to assess ES, many studies focused on a more general landscape scale, rather than specific (grassland) environments. However, the environments in this review were always environments where grassland was present or mentioned, if not the main focus. This shows again that there are relatively few studies that use the concept of ES when assessing attitudes to grassland. This gaps opens up the opportunity for future studies to more explicitly focus on the attitudes to ES in grassland environments, in order to discern the conflicting priorities and to better address these in management practices for sustainability of ES delivery in grassland environments.

Non-farming populations may ascribe more importance to cultural ES over regulating and supporting services due to the fact that the benefits of the latter are less immediate. This might relate to the way in which knowledge about a system is utilised to form attitudes through perceptions, where response to stimuli may be mostly experiential and relate to direct use of a landscape or products rather than a system scale perception of benefits. This may, in turn, be important for understanding how non-farming stakeholder groups appreciate change in a system through their core understanding of personal benefits. Preferences often extended to experiential and local level benefits that would reflect cultural heritage or personal well-being. If such motivations could be more closely incorporated into the delivery of ES at the decision-making and policy scale, there may be better feedback between management practices and the provision of ES that can be beneficial to a diverse set of stakeholders. There is also scope in future research to better understand why there is less positive perception of other types of ES such as regulating and supporting services, particularly in the context of PG environments.

Farmers placed more importance on ES that were connected directly to their farming activity, which is also their immediate experience. Although farmers are more likely to value provisioning and regulating services, cultural services are not unimportant. Farmers’ attitudes to cultural services do not appear to be a direct focus of many studies. Therefore there is potential for the way in which farmers might experience or value cultural ES to be a focus of future studies, for example, there may be cultural connections for farmers based on the longevity of PG sites, some of which can be untouched for generations within a family farm business. The valuation of PG environments may also relate to family values and a sense of stewardship, which studies in this review have shown to be important in forming farmer attitudes.

Studies of conflicting perceptions of ES were rarer, and there is potential for future studies to focus on the trade-offs necessary between stakeholder groups to change systems sustainably. Equally, there were few studies that mentioned attitudes to conflict in relation to ES valuation, which may be an important dynamic of a sustainable system. This would be particularly pertinent for PG landscapes that are under threat, or where new management regimes may be present or likely in the future under changing conditions.

Aesthetics emerged as one of the most important ES. Farmers were found to sometimes hold opposing views to non-farming populations. This contrast means there is potential dissonance in landscapes that are socially acceptable to create or preserve, both for use by stakeholders for recreation and as an element of cultural heritage. There may be implications for changing farmer behaviour, for example, it may be a demotivation if particular management options mean the landscapes becomes visually displeasing to certain farmers under particular management regimes. Intensive farmers were seen to have particular preferences for neat, ordered landscapes, where the land looks economically valuable, perhaps linking to a productivist attitude to farming. Changes to more sustainable management often involve increasing diversity and tend to create ‘messier’ landscapes. Such changes may be barriers for uptake of agro-ecological measures for some farmers (e.g. [120]) Equally, where most people (farmers and non-farmers) often prefer the status-quo, change in landscapes can be difficult to agree upon. The possibility for trade-off and balance of different landscapes is important for meeting sustainability targets, where both types of landscape are likely to be needed in proximity [121]. However, these studies show that where the meaning of the landscape for different stakeholders and groups is taken into consideration in decision-making it is more likely that changes can be better justified, and elements incorporated into new or changing landscapes that continue to hold meaning. Where landscapes are degrading it is also useful to be able to use aesthetic value as a driver for change.

Attitudes to management practice vary amongst groups and between non-farming and farming populations. The underlying perceptions and visions for farming and landscapes often link to resultant attitudes to management options, particularly if the process or the outcomes of the management options do not align with values around the importance of particular ES (e.g. production of food and forage) and factors such as cost. This is an important observation for targeting management interventions. Where attitudes do not match the purpose of the management option uptake and impact will be poor.

Although one study focused on expert stakeholder opinions, missing from the studies were the attitudes of non-farming populations to AES and other management options. A better understanding of the opinions of citizens to the actions and policies applied to landscape management may provide an additional influence on design of management options that are better aligned with demand for ES. Some studies found that the attitudes and perceptions about farming influenced farmers in their uptake of technology as well as the local residents and general population’s attitude to technology in relation to the benefits its application could bring to food quality and safety. As technology becomes increasingly promoted as part of management schemes it is important to understand both farmer and consumer attitudes.

Whilst knowledge is shown to be a key influence on perceptions and attitudes of non-farming and farming stakeholders few studies mentioned knowledge exchange or communication as factors affecting attitudes (in comparison to studies focusing on decision-making and behaviour). It may be a topic for future research to identify links between trust, relationships and attitudes in relation to grassland management. There is also potential to better understand the degree to which personal values can change or are influenced by behaviours or interactions with others, which are not explored deeply in the studies.

Influential factors have been used to profile certain farmer styles (e.g. Hammes et al., 2016), which can be useful to improve understanding and communication among governmental or environmental stakeholders and the farming community. There are few studies that attempt to categorise farmers in this way, perhaps due to the difficulties in containing a diverse population, however the ability to identify patterns in attitudes that link to behaviours is critical to better understanding landscape change and decision-making. Therefore such studies have laid the groundwork for developing flexible and contextual understandings of groups of farmers or non-farmers in relation to PG activities and management.

## 5. Conclusions and implications for PG management

Through a systematic review we show how different stakeholders, including farmers and non-farmers, perceive grassland landscapes and the associated management and ES delivery. Overall, the results of the 135 studies can help to build a picture of the different elements that are important for understanding perceptions and attitudes around PG management, as well as identifying important gaps for future research. Whilst many papers mentioned grassland landscapes, few explicitly focused on attitudes and perceptions to grassland or PG, outside of specific land uses. Research could be better focused on grassland as a land use in relation to attitudes to ES and management in order to better underpin decision-making about PG improvement.

Our analysis shows that key elements of attitudes that can be applied to PG environments include the following:

- understanding how ES are perceived serves as a baseline for the core values that different populations perceive as relevant for farming practices and outcomes;
- attitudes to landscape and landscape change are key to understanding the meaning that different groups see in particular configurations of landscapes, including their aesthetic value; and
- attitudes towards agricultural management help to better understand the behaviours of farmers in relation to uptake, or (in relation to experts or non-farmers) acceptability of interventions.

The study has also revealed factors of importance for influencing attitudes, including:

- contextual factors, which affect underlying experiences, reference points and opportunities of different populations;
- the types of knowledge held by different groups and access to sources of information, as well as the influence of knowledge on behaviour; and
- the role of environmental and personal values in explaining why certain motivations for attitudes and behaviours are held.

As sustainable grassland management requires improvements in the delivery of ES whilst maintaining a profitable farming environment, having an understanding of the attitudes of key decision-makers (farmers), and beneficiaries (non-farmers) is important. When designing and researching new management options there is a need to take into consideration that the basis of farmers’ decision-making is their attitudes, perceptions and beliefs. These are linked to personal circumstances but also to changeable influences such as access to information and policy and economic support. These acknowledgements mean that interventions and changes can be better aligned with farmer attitudes and may have a better uptake. Equally, knowing the attitudes of farmers and non-farmers can mean that much more empowering changes can be implemented. However, we do not yet have a clear solution for adapting the style of agri-environment promotion and engagement to different attitudes and motivations, except that to address conflicting attitudes between farmers and non-farmers it is important to encourage greater integration of these communities, so they understand each other better.

Gaps in this review have been revealed in relation to the need to: review consumer attitudes to grassland landscapes and products; to more deeply understand attitudes to specific ES delivered by grassland landscapes; to research the perceptions of citizens around management of grassland landscapes, and to better understand the drivers of attitudes in relation to contextual factors, knowledge communication and trust. If we can understand the attitudes from both the farmers’ perspective and the publics’ perspective then there is scope to better understand the system as a whole. Each of these groups play either a direct or an indirect role in managing the environments, whether it is making decisions about land use or conservation, or whether it is to value, experience and participate in activities and consume products from grassland environments. This systematic review can inform future exploration of PG landscapes and management from the context of developing sustainable practices in the face of threats and future changes.

---

**S1 Appendix 1. Search strings used in Google Scholar and Scopus.**

**S2 PRISMA Checklist. Details of the systematic review process as reported in manuscript.**

**S3 Protocol. Systematic review protocol.**

